# A role for long-range, through-lattice coupling in microtubule catastrophe

**DOI:** 10.1101/443283

**Authors:** Tae Kim, Luke M. Rice

## Abstract

Microtubules are cylindrical polymers of αβ-tubulin that play critical roles in fundamental processes like chromosome segregation and vesicular transport. Microtubules display dynamic instability, switching stochastically between growing and rapid shrinking as a consequence of GTPase activity in the lattice. The molecular mechanisms behind microtubule catastrophe, the switch from growing to rapid shrinking, remain poorly defined. Indeed, two-state stochastic models that seek to describe microtubule dynamics purely in terms of the biochemical properties of GTP- and GDP-bound αβ-tubulin incorrectly predict the concentration-dependence of microtubule catastrophe. Recent studies provided evidence for three distinct conformations of αβ-tubulin in the lattice that likely correspond to GTP, GDP.P_i_, and GDP. The incommensurate lattices observed for these different conformations raises the possibility that in a mixed nucleotide state lattice, neighboring tubulin dimers might modulate each other’s conformations and hence their biochemistry. We explored whether incorporating a GDP.P_i_ state or the likely effects of conformational accommodation can improve predictions of catastrophe. Adding a GDP.P_i_ intermediate did not improve the model. In contrast, adding neighbor-dependent modulation of tubulin biochemistry improved predictions of catastrophe. Conformational accommodation should propagate beyond nearest-neighbor contacts, and consequently our modeling demonstrates that long-range, through-lattice effects are important determinants of microtubule catastrophe.

## Introduction

Microtubules (MTs) are hollow cylindrical polymers of αβ-tubulin that have essential roles segregating chromosomes during cell division, organizing the cytoplasm, establishing cellular polarity, and more (Desai and Mitchison, 1997). These fundamental activities depend critically on dynamic instability, the stochastic switching of MTs between phases of growing and rapid shrinking (Mitchison and Kirschner, 1984). Dynamic instability is itself a consequence of αβ-tubulin GTPase activity and how it affects interactions between αβ-tubulin in the lattice and at the microtubule end. Although a predictive molecular understanding of catastrophe remains elusive, the broad outlines of an understanding have been established (Mitchison and Kirschner, 1984; VanBuren *et al.*, 2002; Gardner *et al.*, 2011b; Bowne-Anderson *et al.*, 2013; Brouhard, 2015; Duellberg *et al.*, 2016; Brouhard and Rice, 2018). Unpolymerized, GTP-bound αβ-tubulin subunits readily associate at the growing tip of the MTs. Once incorporated into the lattice, αβ-tubulin GTPase activity is accelerated. The assembly-dependence of GTPase activity results in a “stabilizing cap” of GTP- or GDP.P_i_-bound αβ-tubulin near the end of the growing microtubules. Loss of this stabilizing cap triggers catastrophe, the switch from growing to rapid shrinking, because it exposes the more labile GDP-bound microtubule lattice.

Two broad classes of computational models have been developed as part of longstanding efforts to understand in quantitative terms the connections between the properties of individual αβ-tubulins and the polymerization dynamics they collectively generate. ‘Biochemical’ models attempt to recapitulate microtubule dynamics purely in terms of discrete ‘elementary’ molecular reactions like association, dissociation, and GTPase activity (Chen and Hill, 1983; 1985; Bayley *et al.*, 1989; 1990; Martin *et al.*, 1993; VanBuren *et al.*, 2002; Gardner *et al.*, 2011b; Margolin *et al.*, 2012; Piedra *et al.*, 2016) ‘Mechanochemical’ models (Molodtsov *et al.*, 2005; VanBuren *et al.*, 2005; Coombes *et al.*, 2013; Zakharov *et al.*, 2015; McIntosh *et al.*, 2018) use additional spring-like energies to account for conformational strain inside individual αβ-tubulins and for how the resulting mechanical stress affects interactions with other αβ-tubulins in the lattice. A third class of ‘phenomenological’ models (Flyvbjerg *et al.*, 1994; Brun *et al.*, 2009; Bowne-Anderson *et al.*, 2013; Duellberg *et al.*, 2016) uses simplifying assumptions that obscure the relationship between tubulin biochemistry and observable MT behaviors, so we do not consider them further here. Biochemical and mechanical models can each recapitulate microtubule growing and shrinking, and in both kinds of model, catastrophe emerges naturally as a consequence of GTPase activity.

Biochemical models are computationally inexpensive and relatively simple to interpret because they only contain a small number of adjustable parameters. In principle, all these parameters represent measurable quantities that could be accessible to testing/perturbation using site-directed mutagenesis. A limitation of these biochemical models is that they fail to capture the correct concentration dependence and other aspects of catastrophe (e.g. (VanBuren *et al.*, 2002; Bowne-Anderson *et al.*, 2013; Piedra *et al.*, 2016)). The mechanochemical models are computationally expensive and more complicated to interpret because they are more parameter intensive. Some of the parameters describing the spring-like properties of αβ-tubulin might also be hard to validate experimentally. However, the mechanochemical models better recapitulate the concentration-dependence and other aspects of catastrophe where biochemical models fail (Coombes *et al.*, 2013; Zakharov *et al.*, 2015). Why mechanochemical models better capture the concentration-dependence of MT catastrophe remains unclear.

In both biochemical and mechanochemical models, only two nucleotide states are used: GTP and GDP. However, recent structural studies (Alushin *et al.*, 2014; Zhang *et al.*, 2015; Manka and Moores, 2018) have revealed three mutually incommensurate conformations of αβ-tubulin in the body of MT: an ‘expanded’ form that corresponds to an all-GTP lattice, a ‘compacted’ form that correspond to an all-GDP lattice, and an intermediate ‘compact-twisted’ form that correspond to an all-GDP.P_i_ lattice. Because each conformation prefers a different lattice geometry, they must presumably accommodate each other in mixed nucleotide regions of the microtubule lattice. Reconstitution and structural studies of plus-end tracking EB proteins (Maurer *et al.*, 2011; 2012; 2014; Zhang *et al.*, 2015) support a role for these conformations in MT dynamics and regulation. Experiments with a slow shrinking ‘conformation cycle’ mutant of yeast αβ-tubulin (Geyer *et al.*, 2015) that in the GDP state apparently does not relax all the way to the compacted conformation provided evidence that the αβ-tubulin conformation cycle contributes directly to dictate microtubule shrinking rate and catastrophe frequency. It seemed plausible to us that not accounting for a GDP.P_i_ intermediate, or for the likely modulating influence of conformational accommodation in a mixed nucleotide state lattice (Brouhard and Rice, 2018), might explain why biochemical models fail to capture the concentration-dependence of catastrophe.

In the present study, we sought to investigate the consequences of incorporating various candidates for “missing state/biochemistry” into a computational model, with the aim of better predicting the concentration-dependence of catastrophe. We elaborated a Monte Carlo-based algorithm developed in the lab (Ayaz *et al.*, 2014; Piedra *et al.*, 2016; Mickolajczyk *et al.*, 2018) to test if incorporating a GDP.P_i_ state or long-range coupling (reflecting conformational accommodation) improved predictions of microtubule catastrophe. We incorporated the GDP.P_i_ state and conformational coupling separately for simplicity and to be able to assess the effect of each change in isolation. We did not explicitly incorporate ‘mechanochemistry’ into the model because our goal was to identify minimal additions to biochemical models that improve their performance with respect to predicting catastrophe.

Our simulations revealed that incorporating a GDP.P_i_ intermediate state does very little to improve the predicted concentration dependence of catastrophe frequency. Long-range through-lattice conformational accommodation, acting to modulate GTPase rate or dissociation rates, did improve the predictions of catastrophe and its concentration-dependence. Artificially restricting this modulation to short range abrogated the previously observed improvement. Thus, it seems that long-range, through-lattice interactions are important for recapitulating the concentration-dependence of catastrophe in biochemical models. Because mechanochemical models effectively distribute strain throughout the lattice, long-range coupling may represent the specific feature that explains why mechanochemical models have been more successful at predicting catastrophe. By highlighting the importance of long-range, through-lattice effects, our computational experiments provide a new way to think about how catastrophe occurs.

## Results

### A two-state model for microtubule dynamics fails to capture the weak concentration-dependence of catastrophe frequency

We refined our prior algorithm (Ayaz *et al.*, 2014; Piedra *et al.*, 2016; Mickolajczyk *et al.*, 2018) that used kinetic Monte Carlo (Gillespie, 1976; Gibson and Bruck, 2000) to simulate microtubule dynamics. The algorithm simulates one biochemical event (dimer association, dissociation, and GTP hydrolysis) at a time and therefore provides a ‘movie’ of microtubule dynamics. As is commonly done (Chen and Hill, 1985; VanBuren *et al.*, 2002; Molodtsov *et al.*, 2005; Gardner *et al.*, 2011a; Margolin *et al.*, 2012; Zakharov *et al.*, 2015), our model uses a two-dimensional representation of the microtubule lattice to track different kinds of binding environments or neighbor states (Fig. 1A). To minimize the number of adjustable parameters in the model, we initially adopted a very simple parameterization that does not explicitly account for different conformations of αβ-tubulin (reviewed in (Brouhard and Rice, 2014)) and that also does not attempt to describe “mechanical” properties of αβ-tubulin and microtubules such as spring-like conformational strain (reviewed in (Brouhard and Rice, 2018)) (Fig. 1A). The assumptions of this minimalist parameterization are: (i) there are only two nucleotide states (GTP and GDP), (ii) nucleotide is ‘trans-acting’ (Fig. 1A), meaning the strength of the longitudinal interface between dimers (thus the dimer binding affinity at the MT tip) is determined by the nucleotide located at the interface (Rice *et al.*, 2008; Piedra *et al.*, 2016), (iii) the αβ-tubulin dissociation rate for a given subunit determined by the total sum of free energies of all longitudinal and lateral inter-dimer interactions with other subunits, (iv) the association rate into a given site does not depend on the tip configuration, and (v) GTP hydrolysis occurs at the inter-dimer interface, meaning that GTP cannot be hydrolyzed on the most terminal subunit of any protofilament (Fig 1B). In these kinds of models, catastrophe and rescue occur ‘naturally’ (Fig. 1C) in a way that depends on the specific parameters used. Our algorithm is constructed in a highly modular way that makes it easy to implement different biochemical assumptions (Piedra *et al.*, 2016; Mickolajczyk *et al.*, 2018). Later in the paper, we relax the minimalistic assumptions of the two state model to test if more complicated models that incorporate other states or kinds of biochemistry better predict the concentration dependence of catastrophe.

**Figure 1.**
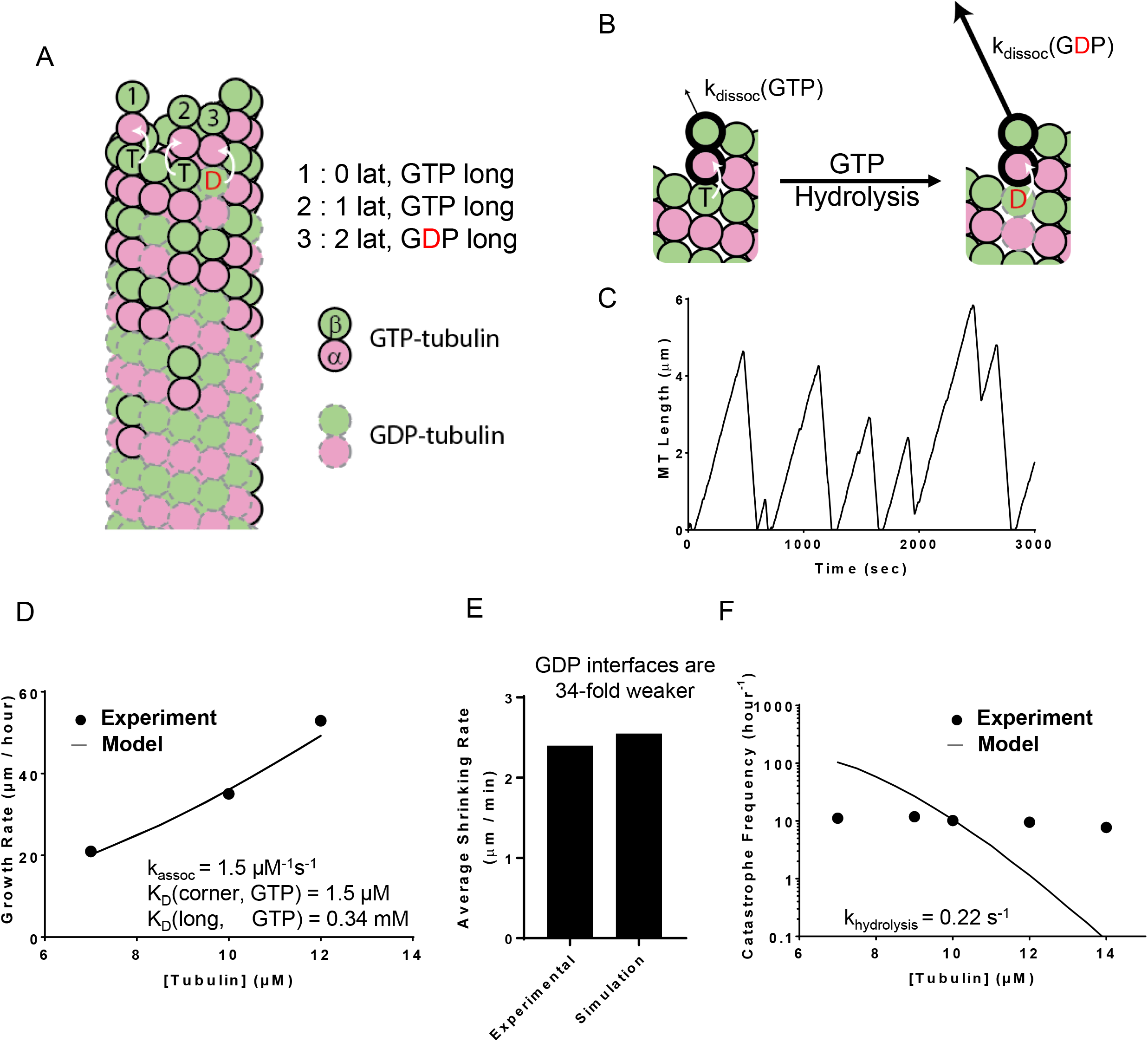
Simulations of a 2-state biochemical model for microtubule dynamics. (**A**) Cartoon representation of a typical growing MT tip during a simulation. αβ-tubulin dimers are represented as pink and green circles; solid black and dashed grey outlines indicate GTP and GDP states, respectively. Dissociation rates depend on the number of lateral neighbors and the identity of the nucleotide at the longitudinal interface (white arrows indicate trans-acting nucleotide, see B). (**B**) Illustration of trans-acting nucleotide. αβ-tubulins with GTP at the longitudinal interface dissociate slower than αβ-tubulins with GDP at the longitudinal interface. (**C**) Representative plot showing simulated MT length vs time at 10 *μ*M αβ-tubulin. The simulation parameters are given in panels D – F. Catastrophes occur naturally as a consequence of the biochemical rules. (**D**) Comparison between measured (black circles) and predicted (line) growth rates. Experimental data are taken from (Gardner *et al.*, 2011b; Coombes *et al.*, 2013). (**E**) Comparison between measured and predicted shrinking rates. (**F**) Comparison between measured (black circles) and predicted (line) catastrophe frequency at different αβ-tubulin concentrations. The 2-state model cannot recapitulate the measured concentration-dependence of the catastrophe frequency.

To obtain model parameters that could recapitulate MT elongation and shrinking rates and approximate the frequency of catastrophe, we followed the divide-and-conquer approach outlined previously (VanBuren *et al.*, 2002; Piedra *et al.*, 2016). We trained our model on recent data that reported growth rates, shrinking rates, and catastrophe frequencies at multiple tubulin concentrations under consistent conditions (Gardner *et al.*, 2011b; Coombes *et al.*, 2013). First, we used “GTP-only” simulations to search for parameters that recapitulated MT growth rates over a range of αβ-tubulin concentrations (Fig. 1D). With those parameters fixed, we optimized the weakening effect of GDP on the longitudinal interface by tuning it to make “all-GDP” microtubules depolymerize at the observed average rate of post-catastrophe shrinking (Fig. 1E). With that new parameter also fixed, we refined the GTPase rate to produce the correct frequency of catastrophe (Fig. 1F). These parameters are not perfectly independent from each other, so we applied this approach iteratively (see Methods). For generality, we also trained our model against ‘classic’ measurements of MT dynamic instability (Walker *et al.*, 1988), where relative to (Gardner *et al.*, 2011b; Coombes *et al.*, 2013) faster shrinking rates and slightly steeper concentration dependence of catastrophe frequency were observed (Supp. Fig. 1). As observed in earlier studies, the predicted catastrophe frequency varies much more strongly with tubulin concentration than observed in experiments (VanBuren *et al.*, 2002; Bowne-Anderson *et al.*, 2013; Piedra *et al.*, 2016). Because the model could not recapitulate the concentration-dependence of catastrophe, we chose 10 *μ*M (the median concentration) as the reference concentration for determining GTPase rate.

### Incorporating a GDP-P_i_ intermediate state into the model does not improve prediction of the concentration dependence of catastrophe

The overly steep concentration dependence of catastrophe predicted by the two-state model may occur because the model does not account for a state or kind of interaction that is important for catastrophe. We added a GDP.P_i_ intermediate between GTP and GDP to test if a three-state model would better predict the concentration dependence of catastrophe. We made the following additional assumptions when implementing the GDP.P_i_ state (Fig. 2A): (i) P_i_ (phosphate) release from the body of the lattice is a first order process, like GTPase, and (ii) the phosphate dissociates instantaneously when exposed at the tip. These new assumptions in the GDP-P_i_ model require two additional parameters: one that describes the strength of a longitudinal contact when GDP.P_i_ is at the interface, and the other that describes the rate of P_i_ release (Fig. 2A).

**Figure 2.**
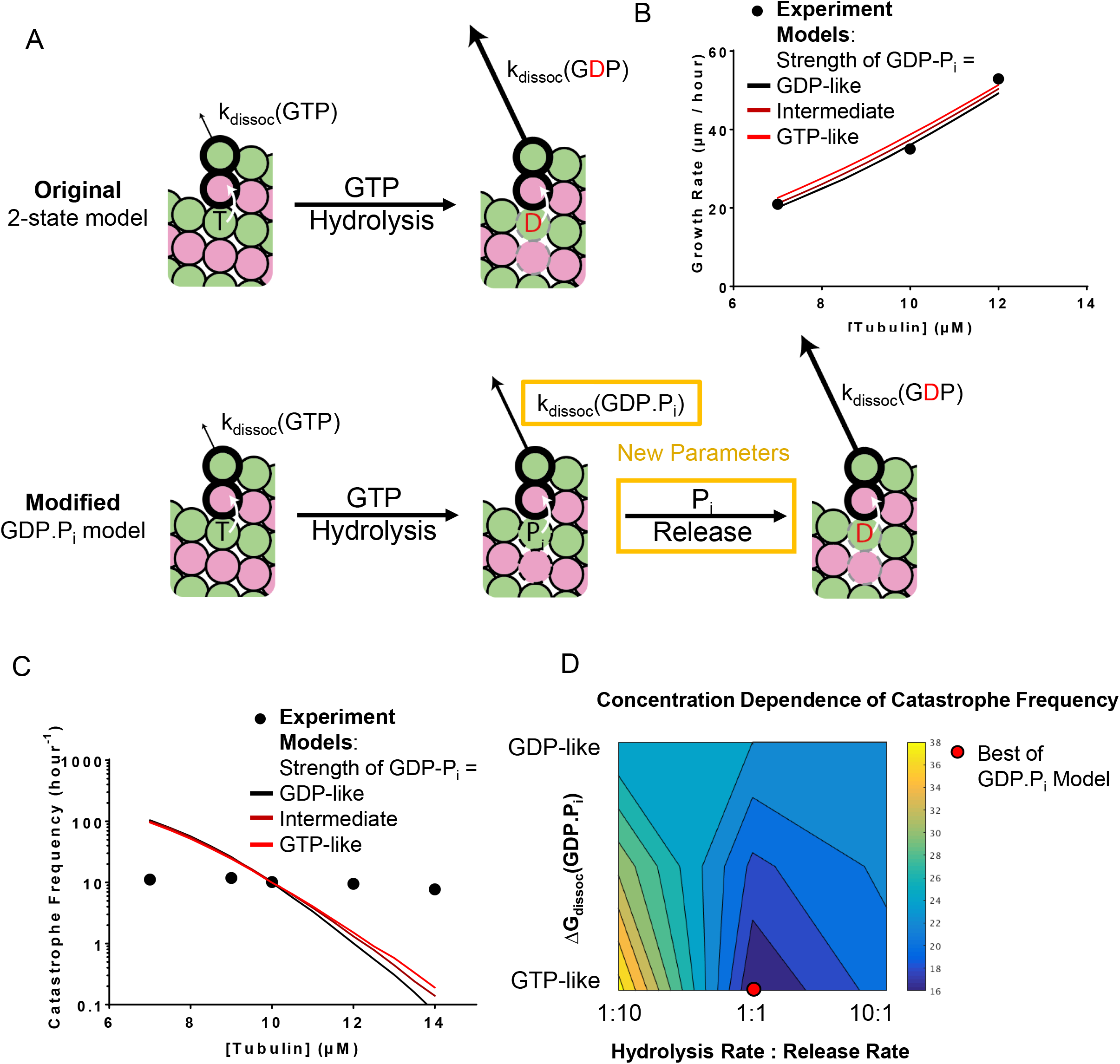
A three state model that contains a GDP.P_i_ intermediate. (**A**) Cartoons illustrating the differences between models without (top) and with (bottom) a GDP.P_i_ intermediate. The GDP.P_i_ intermediate requires two additional parameters: a rate constant for P_i_ release, and another for the strength of the longitudinal interaction when GDP-P_i_ is at the interface. (**B**) Comparison between measured (black circles) and predicted (lines; red, black correspond to GDP.P_i_ interfaces having identical strength as GTP and GDP, interfaces respectively; brown corresponds to GDP.P_i_ interfaces having intermediate strength) growth rates. All three scenarios can recapitulate observed growth rates. In this plot the ratio between the hydrolysis rate and the phosphate release rates have been set to 1:1. (**C**) Predicted catastrophe frequency as a function of concentration for different values for the strength of the GDP.P_i_ longitudinal interface. Varying the strength of the GDP-P_i_ interface has a limited effect on the concentration dependence of the catastrophe frequency. The ratio between the hydrolysis rate and the phosphate release rates have been set to 1:1. (**D**) Contour plot of the predicted concentration dependence of catastrophe. The concentration-dependence is defined as the ratio of catastrophe frequencies at 9 *μ*M and 12 *μ*M. The concentration dependence of the catastrophe frequency is at its lowest when the ratio between the hydrolysis and release is 1:1 and the strength of the longitudinal interface with GDP-P_i_ is as strong as the interface with GTP.

We first examined how varying the strength of longitudinal contacts at the GDP-P_i_ interface affects the predicted frequency of catastrophe as a function of αβ-tubulin concentration. We varied the strength of the GDP-P_i_ interface from strong (equivalent to GTP interface), to intermediate (halfway between GTP and GDP interface), to weak (equivalent to GDP interface), keeping the ratio of the hydrolysis and the release rate constant. Note that setting the strength of the GDP-P_i_ interface to be identical to the GDP interface yields a model that is functionally identical to the two-state model. Whether GTP-like, GDP-like, or in between, the strength of the GDP-P_i_ interface has little effect on predicted growth rates (Fig. 2B). However, and as expected, increasing the strength of GDP.P_i_ interface reduces the catastrophe frequency because it effectively reduces the rate of subunit dissociation from the microtubule end. For a consistent comparison of the concentration dependence of the catastrophe frequency, we retrained the GTPase rate to match the catastrophe frequency at the reference concentration (10 *μ*M) for the strong and intermediate strength GDP.P_i_ interface (while keeping the P_i_ release rates identical to the new GTPase rates, as stated above). The GTPase rate must be increased to compensate for decreased catastrophe frequency (Supp. Table 1A). The newly trained GTPase rates and the strength of the GDP-P_i_ interface, whether GTP-like, GDP-like, or in between, had little effect on the predicted growth rates (Fig. 2B). Keeping the ratio of hydrolysis rate and the release rate same, the predicted concentration dependence of catastrophe frequency does not substantially improve (Fig. 2C).

We then used a grid search approach to explore how changing the ratio between the GTPase rate and the phosphate release rates affects the concentration dependence of catastrophe. We fixed the rate of phosphate release to be 10 times faster or slower than the rate of GTPase and varied the strength of the GDP.P_i_ interface (with re-training of the GTPase rate as described above) as before. In both cases, these changes exacerbated the problems with the model: the predicted concentration-dependence of catastrophe frequency actually increased (Fig. 2D). We observed similar trends in fits to the other dataset that we trained our model against (Walker *et al.*, 1988) (Supp. Fig. 1A). The predicted concentration-dependence of catastrophe was at its lowest when the GTPase rate and the phosphate release rate were the same and when the strength of the GDP.P_i_ interface was as strong as the interface with GTP. However, adding a GDP.P_i_ state did not substantially improve prediction of the concentration-dependence catastrophe.

### Nearest-neighbor conformational accommodation improves predictions of the concentration dependence of catastrophe when modulating GTPase, but not αβ-tubulin dissociation

The expanded conformation (seen in the all GTP lattice) and the compacted conformation (seen in the all GDP lattice) make lattices with different spacing of the lateral interfaces and other changes (Alushin *et al.*, 2014; Zhang *et al.*, 2015; Manka and Moores, 2018). How αβ-tubulins accommodate incommensurate GTP- and GDP-bound conformations in a mixed nucleotide state lattice, as must occur near the tip of the growing MT, is not understood (reviewed in (Brouhard and Rice, 2018)). We speculated that the conformational mismatch might modulate the strength of lateral interactions between αβ-tubulins in different nucleotide states, or that it might modulate the rate of GTPase activity. We implemented these two ideas separately in to the model to test if nearest-neighbor conformational accommodation operating between neighboring αβ-tubulins could improve the predicted concentration-dependence of catastrophe.

To implement neighbor-based modulation of lateral interactions, we assumed that the conformational mismatch/accommodation increases the dissociation rate. In other words, αβ-tubulin with a lateral neighbor that is in a different nucleotide state (and hence conformation) dissociates more quickly than it would otherwise (Fig. 3A). Due to these changes, the ‘nearest-neighbor affinity modulation’ model has only one additional parameter: the fold-faster dissociation rate for αβ-tubulin with a lateral nearest-neighbor with differing nucleotide state. To examine how varying this parameter affects the concentration dependence of catastrophe frequency in simulations, we set the αβ-tubulin with lateral neighbor with different nucleotide to dissociate faster by factors of 1, 1.6 2.7, and 7.8 (these values correspond to free-energy changes of 0, 0.5, 1, and 2 k_B_T, respectively). When the fold increase in dissociation rate is 1, this model behaves identically to the 2-state model. The maximum parameter value of 7.8 is smaller than the GDP weakening factor of 34 for this data set. Further increases in the modulation factor did not lead to substantial reduction in concentration dependence of the predicted catastrophe frequency. We observed similar trends in fits to the other dataset we trained out model against (Walker *et al.*, 1988) (Supp. Fig. 2A). The new parameter only modestly affected the predicted growth rates (Fig. 3B), but at higher values it substantially increased catastrophe frequency. This makes sense, because the exposure of a small number of terminal GDP-bound tubulin can lead to a catastrophe. The GTPase was determined by training the models to the catastrophe frequency at the reference concentration (10 *μ*M), as described above (see also Supp. Table 1B). Compared to the 2-state model, the range of predicted catastrophe frequency over the concentration range decreased from 185-fold to 45-fold (Fig. 3C). Thus, this simple attempt at allowing inter-dimer interaction to be modulated by neighboring nucleotide state somewhat improves the predicted concentration dependence of catastrophe frequency.

**Figure 3.**
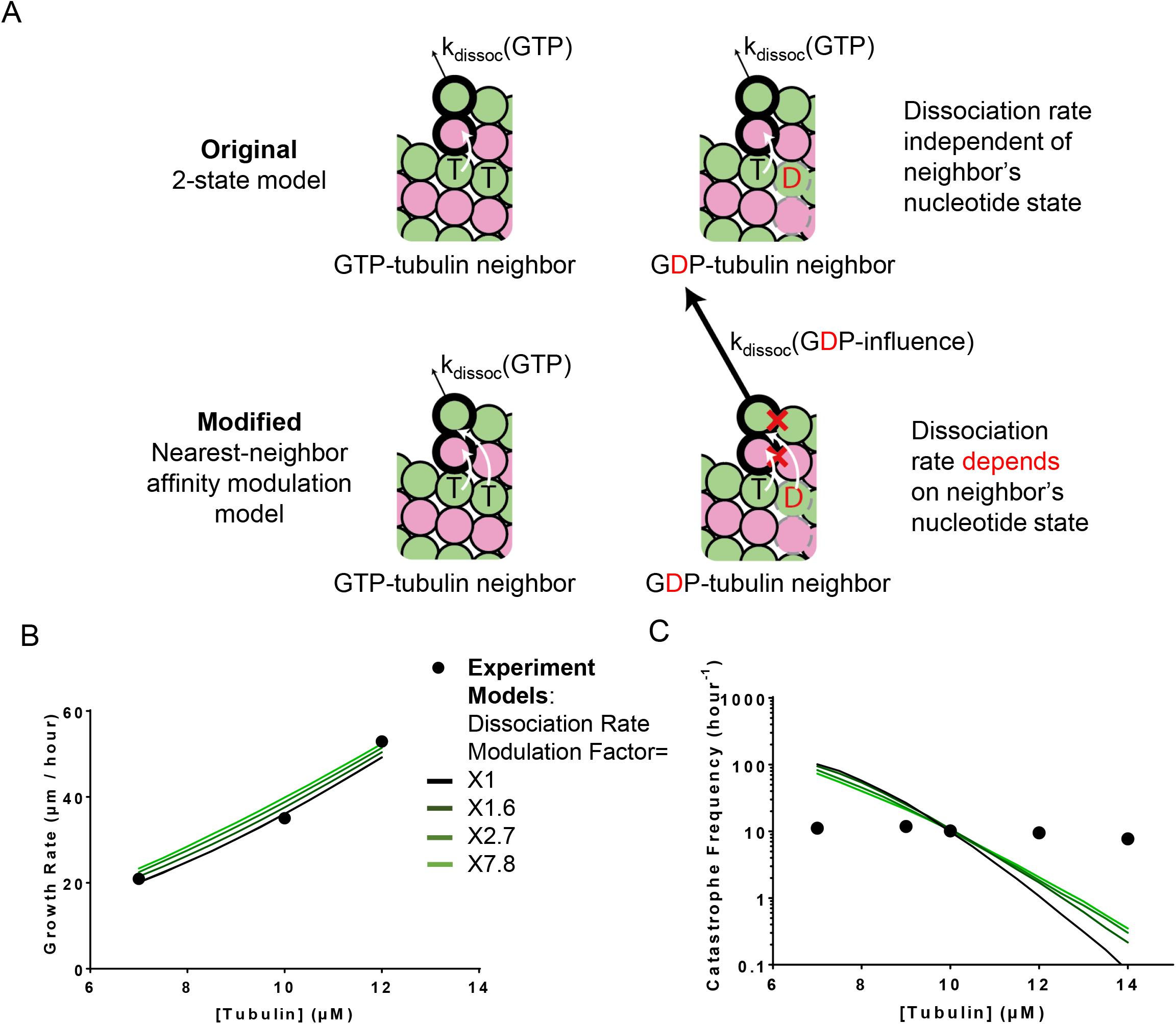
A model that incorporates nearest-neighbor modulation of the strength of lattice contacts. (**A**) Cartoons illustrating the differences between models without (top) and with (bottom) the nearest-neighbor αβ-tubulin affinity modulation. Allowing the αβ-tubulin affinity modulation requires one additional parameter: a fold-increase in αβ-tubulin dissociation rate due to the nearest-neighbor influence. (**B**) Comparison between measured (black circles) and predicted (blackest line corresponds to 1-fold increase in dissociation rates and the greenest corresponds to the 7.8-fold increase) growth rates. All four scenarios can recapitulate observed growth rates. (**C**) Predicted catastrophe frequency as a function of concentration for different fold-increases in αβ-tubulin dissociation rate. Varying the magnitude of αβ-tubulin dissociation modulation has a limited effect on the concentration dependence of the catastrophe frequency.

To implement neighbor-based modulation of GTPase activity, we assumed that αβ-tubulin with GTP next to αβ-tubulin bound to GDP hydrolyzes GTP more quickly (Fig. 4A). In essence, this assumption is equivalent to saying that the ‘accommodating’, intermediate conformation is actually the most active GTPase. This ‘nearest-neighbor GTPase modulation model’ has one additional parameter: the fold increase in hydrolysis rate. We set this neighbor dependent GTPase modulation to increase the rate by factors of 1, 10, 100, and 1000. When the fold increase in hydrolysis rate is 1, the GTPase hydrolysis model is functionally identical to the two-state model. The new parameter did not substantially affect predicted growth rates (Fig. 4B). As before, we adjusted the basal GTPase rate to maintain the correct catastrophe frequency at the reference concentration (Supp. Table 1B). Compared to the two-state model, the range of predicted catastrophe frequency over the concentration range decreased from 185-fold to 7.5-fold. This represents a substantial improvement in the predicted concentration dependence of catastrophe (Fig. 4C). We observed similar trends in fits to the other dataset we trained our model against (Walker *et al.*, 1988) (Supp. Fig. 3A).

**Figure 4.**
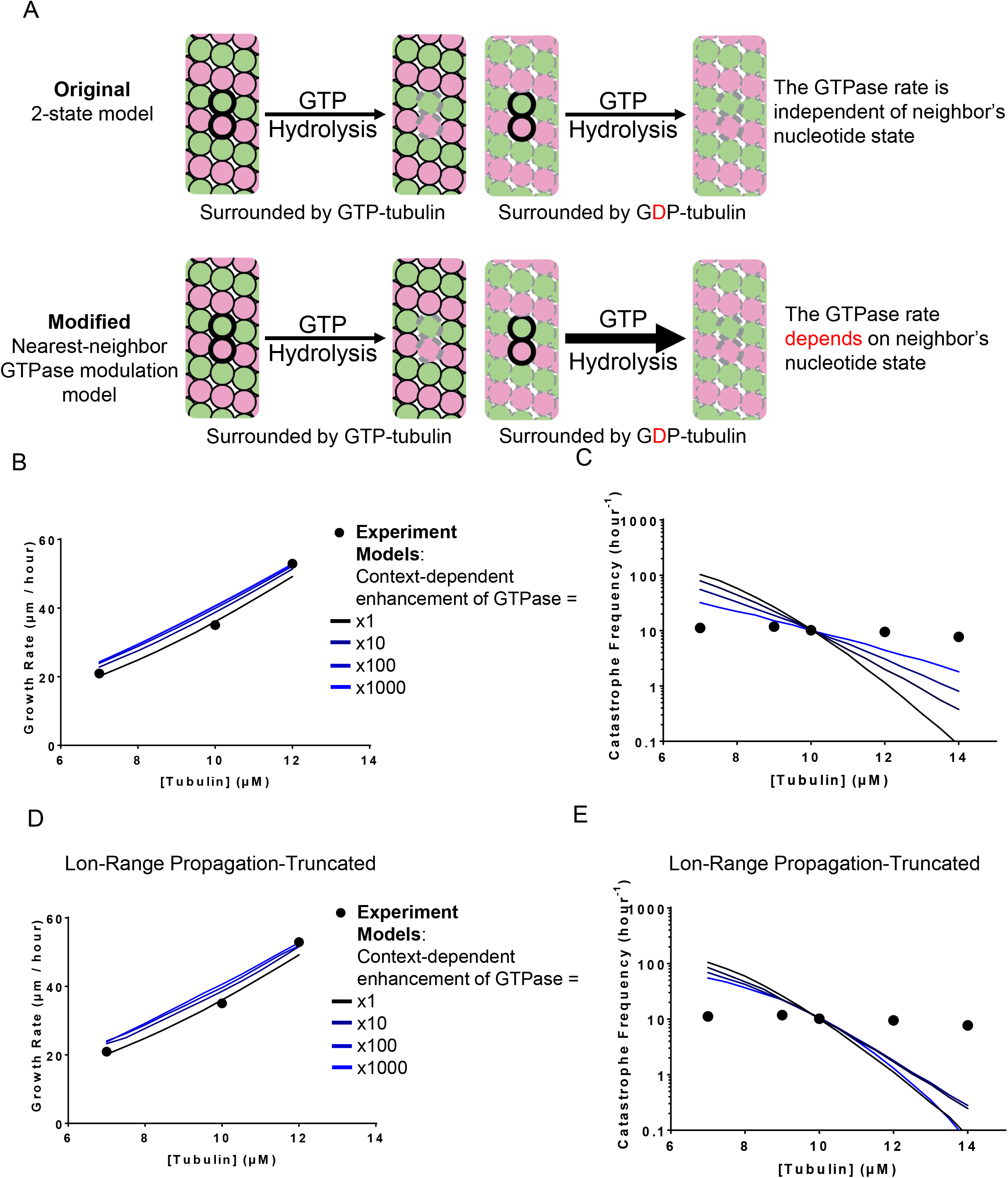
A model that incorporates nearest-neighbor modulation of GTPase activity. (**A**) Cartoons illustrating the differences between models without (top) and with (bottom) the nearest-neighbor GTPase modulation. Allowing the GTPase rate modulation requires one additional parameter: the fold-increase in GTPase rate due to the nearest-neighbor influence. (**B**) Comparison between measured (black circles) and predicted (blackest line corresponds to 1-fold increase in GTPase rates and the bluest corresponds to the 1000-fold increase) growth rates. All four scenarios can recapitulate observed growth rates. (**C**) Predicted catastrophe frequency as a function of concentration for different fold-increases in GTPase rate. Varying the magnitude of GTPase rate modulation has a significant effect on the concentration dependence of the catastrophe frequency. (**D**) Comparison between measured (black circles) and predicted (blackest line corresponds to 1-fold increase in GTPase rates and the bluest corresponds to the 1000-fold increase) growth rates, in the propagation-limited GTPase model. All four scenarios can recapitulate observed growth rates. (**E**) Predicted catastrophe frequency as a function of concentration for different fold-increases in GTPase rate, in the propagation-limited GTPase model. Artificially limiting the propagation of wave-like GTPase activity reverts the changes in predicted concentration dependence of catastrophe frequency observed in the original nearest-neighbor GTPase modulation model.

Why did this nearest-neighbor GTPase modulation model improve predictions of the concentration dependence of catastrophe so much more dramatically? Looking at the biochemical “movies” generated by the simulation, and even though we implemented this as a nearest-neighbor modulation, we observed that the GTP hydrolysis propagated through the lattice, like a wave (not shown). The wave of GTP hydrolysis starts from a random GTP hydrolysis in locally all-GTP lattice, where hydrolysis is relatively slow in this model. Hydrolysis of one GTP to GDP effectively starts a chain reaction because the nearest neighbor αβ-tubulins have increased GTPase activity. GTP hydrolysis at this second site then primes its neighbor for increased GTPase activity, and so on. Thus, although we constructed the model to have only nearest neighbor effects, the resulting behavior showed longer-range propagation.

Was it the local change in GTPase rate or the longer-range propagation of GTP hydrolysis that was most important for improving the predicted concentration-dependence of catastrophe? To examine this question, we modified the nearest-neighbor GTPase modulation model so that the wave of GTP hydrolysis would be artificially prevented from propagating too far. To accomplish this, we disallowed “across-seam” interactions from modulating GTPase activity. As before, we set the neighbor dependent GTPase modulation to increase by factors of 1, 10, 100, and 1000, and retrained the basal GTPase rate to match the catastrophe frequency at the reference concentration. Limiting the propagation of (‘truncating’) the nearest-neighbor stimulation of GTPase degraded the model’s ability to predict the concentration-dependence of catastrophe. Indeed, whereas at low GTPase modulation factors we observed a modest improvement in the predicted concentration-dependence of catastrophe, at higher modulation factors this trend reversed (Fig. 4E). Whereas the untruncated model showed only a 7.5-fold change in catastrophe frequency over the measured concentration range, the truncated version showed a 110-fold change, nearly reverting back to 185-fold change observed in the two-state model. We observed similar trends in fits to the other dataset we trained our model against (Walker *et al.*, 1988), but the magnitude of the difference was much smaller than in the models trained to the data from (Gardner *et al.*, 2011b; Coombes *et al.*, 2013) (Supp. Fig. 3B). In summary, nearest-neighbor modulation of αβ-tubulin dissociation rate had limited effect on the predicted concentration dependence of catastrophe. By contrast, nearest-neighbor modulation of GTPase activity yielded a substantial improvement. The activation of GTPase propagated through the lattice, and we showed that this long-range propagation was required for improved predictions of catastrophe.

### Incorporating long-range through lattice modulation of the strength of tubulin-tubulin interactions can also improve predictions of catastrophe

In the nearest-neighbor GTPase modulation model, the wave-like propagation of GTP hydrolysis effectively allowed the nucleotide state at one site to indirectly affect the biochemistry of distant (beyond nearest-neighbor) αβ-tubulins. We wondered if incorporating long-range through-lattice modulation of αβ-tubulin:αβ-tubulin binding affinity could also improve the predicted concentration dependence of catastrophe.

Previously, in the nearest-neighbor affinity modulation model, for simplicity we assumed that the destabilizing inter-dimer interaction was limited to nearest neighbors. However, it stands to reason that if one subunit influences the conformation of its neighbor, then that neighbor should influence the conformation of its neighbor, and so on. In other words, the conformational accommodation should propagate beyond nearest neighbor contacts. We implemented a version of this model wherein the accommodation modulates the strength of lattice contacts over some specified distance (number of αβ-tubulin subunits), by modifying the affinity model so that the nucleotide state of the tubulins affects the dissociation rate of other tubulins further away. This time, we kept the modulated dissociation rate to 7.4, the highest value tried for the original nearest-neighbor affinity modulation model. Then we varied the maximum range of modulation (Fig. 5A). When the range is set to 0, the model is identical to the two-state model, while if the range is set to 1, the model is identical to the original nearest-neighbor affinity model. When the range is an integer greater than one, it means a given subunit affects that many of its neighbors to the left and to the right. Then, we retrained the GTPase rate to match the catastrophe frequency at the reference concentration. As we increased the maximum range of through-lattice modulation of inter-dimer interaction, the predicted catastrophe frequency became substantially less sensitive to αβ-tubulin concentration (Fig. 5C). Compared to the nearest neighbor model, allowing long-range effects yielded an additional ~4.5-fold decrease in the range of catastrophe frequencies over the concentration range. We observed similar trends in fits to the other dataset we trained our model against (Walker *et al.*, 1988) (Supp. Fig. 2B).

**Figure 5.**
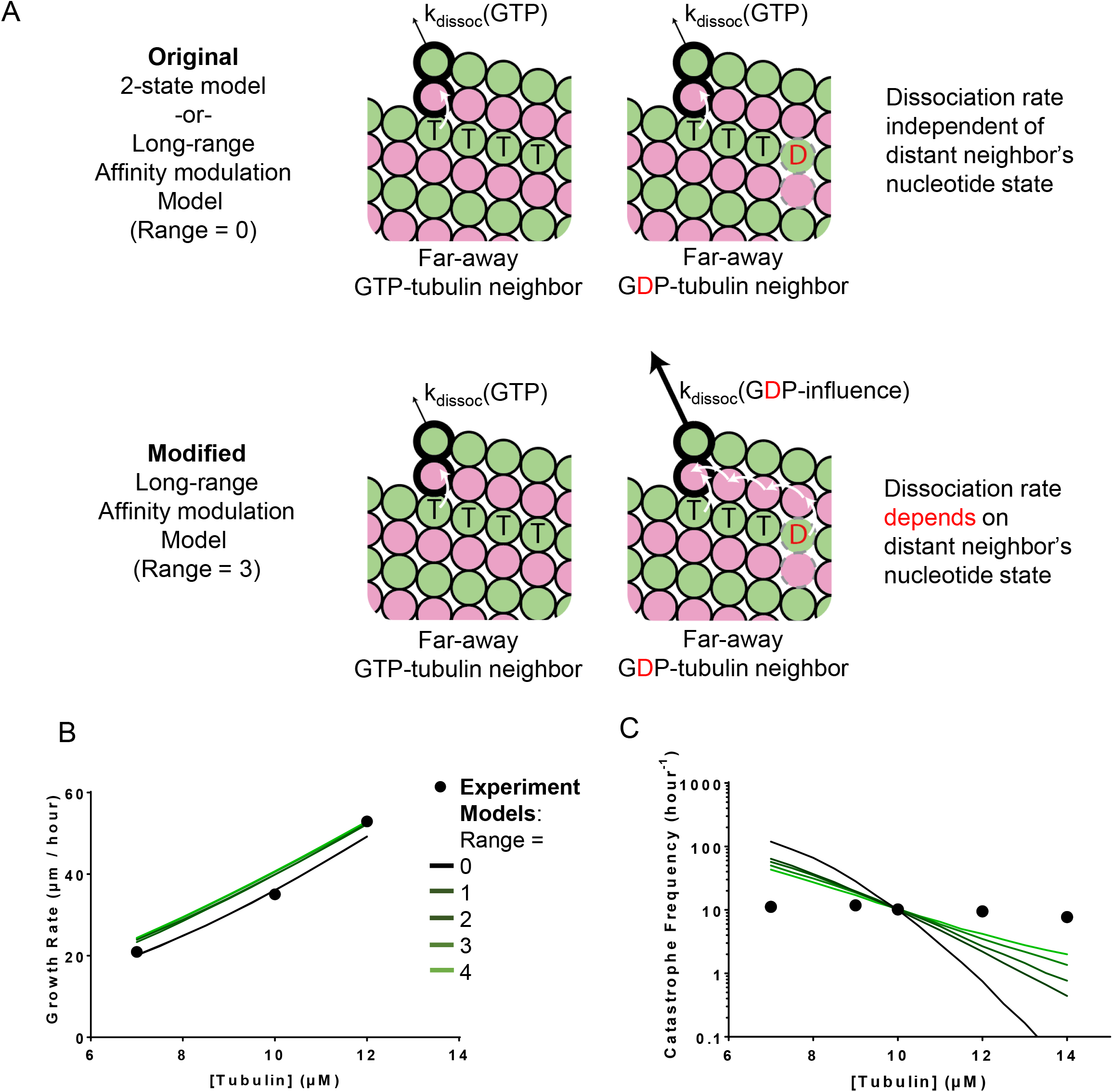
A model that incorporates long-range modulation of the strength of lattice contacts. (**A**) Cartoons illustrating the differences between models without (top) and with (bottom) the long-range αβ-tubulin affinity modulation. Allowing the αβ-tubulin affinity modulation requires two additional parameter: a fold-increase in αβ-tubulin dissociation rate due to the nearest-neighbor influence and the maximum range of modulation. (**B**) Comparison between measured (black circles) and predicted (blackest line corresponds to the modulation range of 0 and the greenest corresponds to the modulation range of 4) growth rates. All five scenarios can recapitulate observed growth rates. In this plot the dissociation rate of the modulated αβ-tubulin is increased by 7.8-fold. (**C**) Predicted catastrophe frequency as a function of concentration for different maximum range of modulation. Varying the maximum range of modulation has significant effect on the concentration dependence of the predicted catastrophe frequency. The dissociation rate of the modulated αβ-tubulin is increased by 7.8-fold.

In summary, incorporating long-range, through-lattice modulation of tubulin-tubulin interactions improved predictions of the concentration-dependence of catastrophe. Short-range modulation was much less effective. Incorporating a ‘third state’ in the form of GDP.P_i_ also did not improve predictions of the concentration-dependence of catastrophe. Thus, it appears that long-range through-lattice effects, whether modulating GTPase or αβ-tubulin dissociation, represent a missing ingredient required for biochemical models to recapitulate the concentration-dependence of catastrophe.

### An empirical decomposition of catastrophe frequency reveals differences in frequency of pausing and commitment to catastrophe between the models

To better understand why incorporating through-lattice long-range modulation improved predictions of the concentration dependence of catastrophe, we took a closer look at the sequence of events that led to catastrophe in our different models. In all models, MT growth always paused (defined as a transient growth rate less than 25% of the average MT elongation rate) for a short time before undergoing a catastrophe (Fig. 6A). Similar pausing/slowdown has been observed in experiments (Maurer *et al.*, 2014). As we showed previously (Piedra *et al.*, 2016), terminal GDP exposure can cause this slowing of elongation by transiently poisoning individual protofilaments. This transient pausing in turn accelerates erosion of the stabilizing cap, and the consequent complete loss of the cap leads to catastrophe. However, not all pausing episodes led to a catastrophe in our simulations. If the GDP exposure can be overcome before the stabilizing cap completely erodes, the MT can resume growing at a normal rate. If the transient pausing is truly an obligate intermediate step between growth and catastrophe, then the product of “growth-to-pause” frequency and “pause-to-catastrophe” probability (not frequency because this is just a score of how catastrophe occurs as a result of transient pausing) should yield the catastrophe frequency (see Methods) (Fig. 6B). We quantified the “growth-to-pause” frequency and the “pause-to-catastrophe” probability from simulation outputs of the 2-state model; their product faithfully reproduced the frequency of catastrophe (Fig. 6C). Thus, transient pausing is necessary but not sufficient for catastrophe, and we can decompose catastrophe into two separate steps.

**Figure 6.**
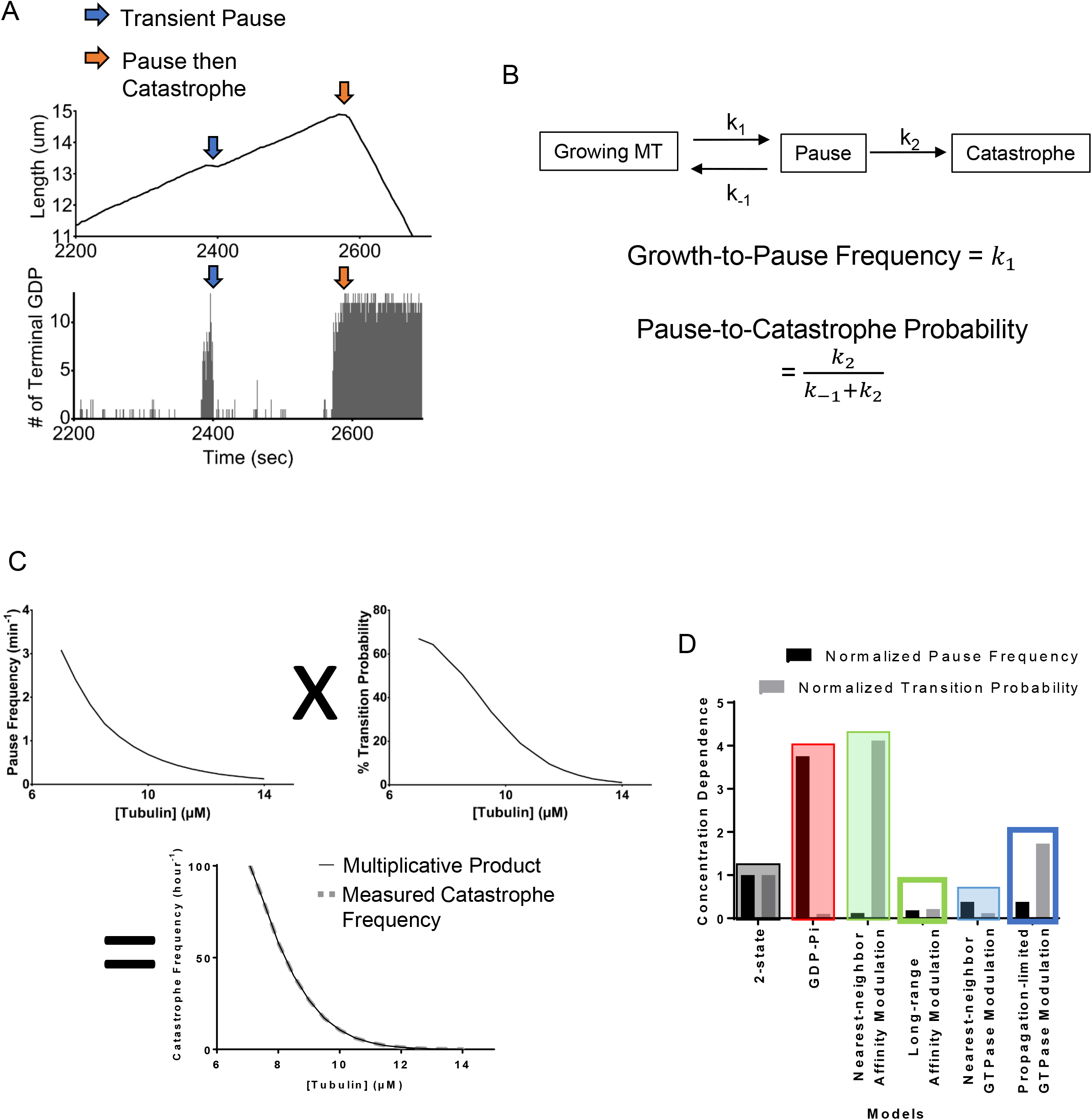
Microtubule catastrophe can be decomposed into two separate steps. (**A**) The plot of the MT length vs time (top panel) and the corresponding plot of terminal GDP-tubulin vs time (bottom panel). The exposure of GDP-tubulins at the end of some protofilaments (blue arrow) leads to a transient pausing. The exposure at the end of all protofilaments can follow partial loss of the GTP stabilizing cap (orange arrow) leading to a transient pausing followed by a catastrophe. (**B**) A diagram of transient pausing and catastrophe as elementary processes (top). “Growth-to-pause” frequencies and “pause-to-catastrophe” probability defined in terms of the reaction rates to the elementary processes (bottom). (**C**) The plot of “growth-to-pause” frequencies (top left), “pause-to-catastrophe” probabilities (top right), and the catastrophe frequencies (bottom) as functions of αβ-tubulin concentrations in two-state model. The multiplicative product (black line, bottom plot) of the “growth-to-pause” frequencies and the “pause-to-catastrophe” probabilities match the value of the predicted catastrophe frequency (gray dashed line, bottom plot). (**D**) The concentration dependencies of the “growth-to-pause” frequencies and the “pause-to-catastrophe” probabilities of different models normalized to two-state model. Here, we defined the concentration dependence as the ratio of the “growth-to-pause” frequencies or the “pause-to-catastrophe” probabilities at 9 *μ*M over 12 *μ*M.

If the catastrophe frequency is the product of the growth-to-pause frequency and the pause-to-catastrophe probability, then the concentration dependence of the catastrophe frequency must also stem from the concentration dependence of its components. To determine if the concentration dependence can be attributed to the growth-to-pause frequency, the pause-to-catastrophe probability, or both, we first measured the growth-to-pause frequency and the pause-to-catastrophe probability as functions of tubulin concentration in the two-state model. Both the growth-to-pause frequency and the pause-to-catastrophe probability depended strongly on αβ-tubulin concentration (Fig. 6C).

We then examined how different models affected the concentration dependencies of growth-to-pause frequencies and pause-to-catastrophe probabilities relative to the baseline provided by the two-state model (Fig. 6D). Compared to the two-state model, all models showed substantial changes to the concentration dependence of growth-to-pause frequency and pause-to-catastrophe probability. In the models that did not allow long-range effects, the concentration-dependence of the two components of catastrophe moved in opposite directions, effectively cancelling each other so that there was little improvement in the concentration-dependence of the catastrophe. By contrast, in the models that allowed long-range effects, the concentration-dependence of the two components of catastrophe moved in concert with each other, explaining why these long-range models better predicted catastrophe.

## Discussion

Simple two-state biochemical models fail to predict the weak concentration-dependence of the catastrophe frequency. The studies described here were motivated by the hypothesis that this failure occurs because two-state models oversimplify the biochemistry, and that we might be able to gain insight into what was missing using modeling. We sought to test whether adding different candidates for missing states or kinds of interactions to a biochemical model for microtubule dynamics could improve predictions about catastrophe. The new kinds of biochemistry we tested were inspired by recent structural experiments (Alushin *et al.*, 2014; Zhang *et al.*, 2015; Manka and Moores, 2018) that revealed three distinct and apparently mutually incommensurate conformations of αβ-tubulin in the GTP, GDP.P_i_, and GDP-bound microtubule lattice. These structural findings, along with results from reconstitution studies of EB proteins (Maurer *et al.*, 2011; 2014), imply that the models might need to contain a third state (GDP.P_i_), or they might need to account for the likely effects of incommensurate conformations αβ-tubulin by modulating the properties of GTP- or GDP-tubulin in a context-dependent way (conformational coupling). A third state and conformational coupling might simultaneously be required, but for simplicity in this work we chose to examine the third state and conformational coupling models separately.

Adding a third state did little if anything to improve predictions of catastrophe. By contrast, allowing conformational coupling to modulate either the GTPase rate or the lattice-binding affinity of terminal subunits noticeably improved predictions of the concentration-dependence of catastrophe. Because this conformational coupling should propagate beyond nearest-neighbor interactions, our computational findings suggest that through-lattice cooperative effects are important determinants of microtubule catastrophe. None of the models we examined fully capture the concentration-dependence of microtubule catastrophe measured in experiments. This should not be surprising, because we intentionally chose the simplest (least parameter intensive) ways to examine the possible consequences of candidate ‘missing biochemistries’ like a GDP.P_i_ state or the coupling that arises from conformational accommodation.

Mechanochemical models have outperformed biochemical models where catastrophe is concerned: mechanochemical models better recapitulate both the concentration- and age-dependence of microtubule catastrophe (Coombes *et al.*, 2013; Zakharov *et al.*, 2015). The mechanochemical models are more parameter intensive, however, and they account for multiple features that the biochemical models do not: curved-straight conformational changes on the microtubule end, long-range energetic coupling in the lattice, longitudinal inter-dimer twist, and more. Consequently, precisely why these mechanochemical models better predict the concentration dependence of catastrophe compared to biochemical models has so far not been clear. The work described here may provide insight into why mechanochemical models have been more successful at predicting catastrophe. Indeed, our simulations indicate that long-range, through-lattice coupling is required for improved predictions of catastrophe in biochemical models. Because of the way that they allow mechanical strain to be distributed through the lattice, a kind of long-range coupling is included in mechanochemical models. In light of our results, it seems likely that the success of mechanochemical models can be attributed to the fact that they incorporate long-range coupling in the lattice.

What we have described based on our modeling is a kind of cooperativity that operates within the microtubule. This resonates with a view of microtubule dynamics (Kueh and Mitchison, 2009) (Brouhard and Rice, 2018) in which different conformations of αβ-tubulin can modulate or even override nucleotide state in dictating biochemical interactions and rates in the lattice. Detecting such cooperativity experimentally and determining whether it operates on GTPase or the strength of lattice contacts are important challenges for future work. The recently introduced ability to work with tubulins from different species (Widlund *et al.*, 2012; Chaaban *et al.*, 2018) (Drummond *et al.*, 2011), to purify single isotypes and site-directed mutants (Johnson *et al.*, 2011; Minoura *et al.*, 2013; Geyer *et al.*, 2015; Pamula *et al.*, 2016; Ti *et al.*, 2016; Vemu *et al.*, 2016; 2017; Geyer *et al.*, 2018) (Drummond *et al.*, 2011), and to measure αβ-tubulin : microtubule interactions at the single molecule level (Mickolajczyk *et al.*, 2018) have the potential to accomplish this, and promise to provide new kinds of data that will drive a deeper understanding of microtubule catastrophe.

In summary, our computational experiments demonstrate that beyond-nearest-neighbor, through-lattice effects can make important contributions to microtubule catastrophe. The combination of this allosteric conformational coupling with the extended microtubule lattice has the potential to generate abrupt, switch-like changes (reviewed in (Bray, 2013) for other systems) that could give rise to threshold-type behaviors wherein the switch only happens upon reaching some critical percentage of GTP-hydrolysis (or some other property). Interestingly, the onset of rapid shrinking has been observed to occur after exceeding a threshold loss of the stabilizing cap (Maurer *et al.*, 2014). A number of microtubule-associated proteins have recently been shown to alter the microtubule lattice upon binding (Zhang *et al.*, 2015; Shima *et al.*, 2018) (Zhang *et al.*, 2017) (Howes *et al.*, 2017) (Loeffelholz *et al.*, 2017; Peet *et al.*, 2018; Zhang *et al.*, 2018), and these binding-induced conformational changes might also modulate properties of the lattice at greater distance. At least one study has proposed that EB proteins might influence the activity of XMAP215-family microtubule polymerases via long-range, through-lattice effects (Zanic *et al.*, 2013), but the underlying mechanism was not specified. The apparent importance of long-range cooperative/allosteric effects suggests that material-like properties of the microtubule are important for catastrophe and may be targeted by regulatory factors.

## Acknowledgements

We thank B. Geyer, S. Majumdar, E. Bonventre, and X. Ye for critical comments on the manuscript. LMR is the Thomas O Hicks Scholar in Medical Research. This work was supported by a grant from the NSF to LMR (MCB-1615938). TK received support from NIH T32 GM008297.

## Methods

### Computational simulation of the models

We created a computer program (coded in fortran) to perform kinetic Monte Carlo simulations of MT plus ends. The model is similar to one we used previously (Ayaz *et al.*, 2014; Piedra *et al.*, 2016; Mickolajczyk *et al.*, 2018), and was inspired by an earlier implementation from others (VanBuren *et al.*, 2002). Briefly, the microtubule lattice is represented by a two dimensional array with a periodic boundary condition to mimic the cylindrical wall of the microtubule. MT dynamics is simulated one biochemical reaction (αβ-tubulin subunit association or dissociation, and GTP hydrolysis) at a time. In a prior study we reported that the rate of GDP to GTP exchange on the microtubule end could modulate the frequency of catastrophe (Piedra *et al.*, 2016). That reaction did not improve the predicted concentration-dependence of catastrophe (Piedra *et al.*, 2016), so for simplicity we did not include it in the models described here. For the two-state model, the association can happen at the tip of each protofilament, and association rate is given by k_on_ × [αβ-tubulin], where k_on_ denotes the on rate constant. The terminal subunits can dissociate from the MT lattice at a rate given by k_on_ × K_D_, where K_D_ is the affinity determined by the sum of all inter-dimer interactions. As described previously (Piedra *et al.*, 2016), our parameterization assumes that the nucleotide (GTP or GDP) acts in-trans to affect the strength of longitudinal contacts such that GTP contacts are stronger than GDP ones. GTP hydrolysis is modeled for all nonterminal subunits with rate constant k_hyd_.

### Automated analysis of simulations

We created custom MATLAB routines to analyze the output from the simulations. These routines determine the instantaneous growth / shrinking rates by looking at the change in the total number of subunits over a 5 second time period. If the instantaneous growth rate falls below 25% of the average growth rate during the growth phase, the simulated MTs are considered to have paused for the duration of the slower growth. The pause episodes are left out of the growth / shrinking rate calculations and are used to determine how frequently the simulation pauses. The MATLAB routine automatically detects MT catastrophe using the following definition: the simulated MT persistently must be shrinking at a rapid rate (shrinking rate greater than 75% of the mode of shrinking rate distribution) for an extended period of time (at least 15 seconds). In the two-step decomposition of catastrophe, the frequency of pausing is tabulated to obtain the ‘growth to pause’ frequency, and the likelihood of catastrophe following transient pausing gives the “pause-to-catastrophe” probability. We used the ratios between values at 12 *μ*M and 9 *μ*M as a measurement of the concentration dependencies of the “growth-to-pause” frequency and the “pause-to-catastrophe” probability. These ratios (the concentration dependencies) were normalized to the concentration decencies of the two-state models for model to model comparisons.

### The parametrization of two-state computational model

To parametrize the two-state model, we first assumed a value for k_on_ (1.5 tubulin·s^−1^·*μ*M^−1^ per protofilament for the data that we fit in the main text (Gardner *et al.*, 2011b; Coombes *et al.*, 2013) and 2 tubulin·s^−1^·*μ*M^−1^ per protofilament for the alternative data that we fit in the supplemental material (Walker *et al.*, 1988)). Then, the strengths of the longitudinal (at the GTP-interface) and lateral interaction were determined by fitting the model predictions on growth rates (during the growing phase) to the experimental values. The strength of the longitudinal interaction at the GDP-interface was determined by fitting the model predictions on shrinking rates (during the shrinking phase) to the experimental values. Then the GTPase rate is determined by fitting the model predictions on the catastrophe frequency at a single reference concentration (10 *μ*M for the data (Gardner *et al.*, 2011b; Coombes *et al.*, 2013) in the main text and 12 *μ*M for the alternative set (Walker *et al.*, 1988) in supplemental material). This process is repeated until all adjustable parameters stabilize.

### The GDP.P_i_ model

We incorporated a third intermediate state into our model, by including GDP.P_i_. This ‘GDP.P_i_ model’ inherits the two-state model’s properties described above with some modifications. In this model, GTP is first hydrolyzed to GDP.P_i_ then the P_i_ released at a set rate to form GDP. We assumed that the P_i_ is released immediately when exposed at the tip of the MT, and that the strength of the longitudinal interface with GDP.P_i_ is different from the ones with GTP or GDP. This model has two additional parameters: the rate of P_i_ release and the strength of the longitudinal interface with GDP.P_i_. We explored the parameter space of the additional adjustable parameters in a 3-by-3 grid pattern: setting the rate of P_i_ release 0.1, 1, 10-fold of the GTPase rate, and setting the strength of the longitudinal interface with GDP.P_i_ to strong (GTP-like), intermediate, and weak (GDP-like). Then, in order to maintain the correct frequency of catastrophe at the reference concentration, we retrained the GTPase rate. We kept the kept the αβ-tubulin binding affinities the same as in the two-state model because we did not want to introduce confounding variation. Changes in growth and shrinking rates due to the modification were negligible.

### The affinity modulation models

As before, the affinity modulation models inherit the two-state model’s properties described above with some modifications. In the nearest-neighbor affinity modulation model, we assumed that the rate of αβ-tubulin dissociation is faster if the nucleotide state of the longitudinal interface of the nearest neighbor is different. This model has one new adjustable parameter: the energy cost of being next to a tubulin with different nucleotide. We explored the parameter space by setting the energy cost to different values and retraining the GTPase. As described above, we maintained the αβ-tubulin binding affinities the same as in the two-state model. In the long-range affinity modulation model, the range of influence for the affinity modulation is an additional adjustable parameter. When the range is set to 0, the model behaves identically as the two-state model, and when the range is set to 1, the model behaves identically as the nearest-neighbor affinity modulation model; for values larger than 1 it gives beyond-nearest-neighbor effects. For this model, we set the energy cost to the maximum value we used for the nearest-neighbor affinity modulation model and varied the range from 0 to 4.

### The GTPase modulation models

In the nearest-neighbor GTPase modulation model the αβ-tubulin with GTP laterally next to αβ-tubulin bound to GDP hydrolyzes GTP faster. This model has one additional parameter: the context-dependent fold-increase in GTPase rate. We set the fold increase to 1, 10, 100, and 1000, and retrained the basal GTPase rate, as before. This context-dependent increase in GTPase rate leads to a wave-like propagation of GTP hydrolysis. In the propagation-limited GTPase modulation model, we limited the wave-like propagation of the GTP hydrolysis by preventing GTPase modulation across the MT seam.

